## supplementary table and figures for "Linking Molecular Mechanisms to their Evolutionary Consequences: a primer"

**Supplementary tables:**

**Table S1. Sequences of 9 promoters used to validate the model**. The first sequence is the wild-type $P_R$ promoter used in this study. Note that we introduced a varied number of mutations into each promoter in order to cover a wider range of steady-state and dynamical phenotypes: mutants with multi-dimensional phenotypes similar to the wild-type, and those with altered ON, OFF or both ON and OFF steady-state expression levels. The measured gene expression dynamics shown in Figure 2 – Extended Figure 1 show the range of dynamical phenotypes covered by this set of mutants.

| Sequence |
|---|
| GATAAATATTTATCTCTGGCGGTGTTGACATAAATACCACTGGCGGTGATACTGAGCACATCAGCAG |
| GATAAATATTTATATCTGGCGGTGTTGACATAAATACCACTGGCGGTGATACTGAGCACATCAGCAG |
| GATAAATATTTATCTCTGGCGGTGTTGACATAAATACCACTGACGCTGATACTGAGCACATCAGCAG |
| GATAAATATTTATCTCTGGCGGTGTTGACATAAATACCACTGACGCTGATACTGAGCACATCAGCAG |
| GATAAATATTTATCTCTGGCGATGTTGACATAAATACCACTGGCGGTCATACTGAGCACATCAGCAG |
| GATAAATATTTACCTCTGGCGGTGTTGACCTAAATACCACTGGCGGTGATACTGAGCACATCAGCAG |
| GATAAATATTTATCTCTGGCGGTGTTGTCATAAACACCACTGGCGGTGATACTGAGCACATCAGCAG |
| GATAAATATTTATCTCTGGCGGTGTTGACCTAAATACCACTGGCGGTGATACTGAGCACATCAGCAG |
| GATAAATATTTTTCTCTGGCGGTGTTGACATGAATACGACTGGCGGTGATTCTGAGCACATCAGCAG |
| GATAAATATTTATCTCTGGCGGTGTTGACATAAATACCGCTGGCCGTGACACTGAGCACATCAGCAG |

**Supplementary Figures:**

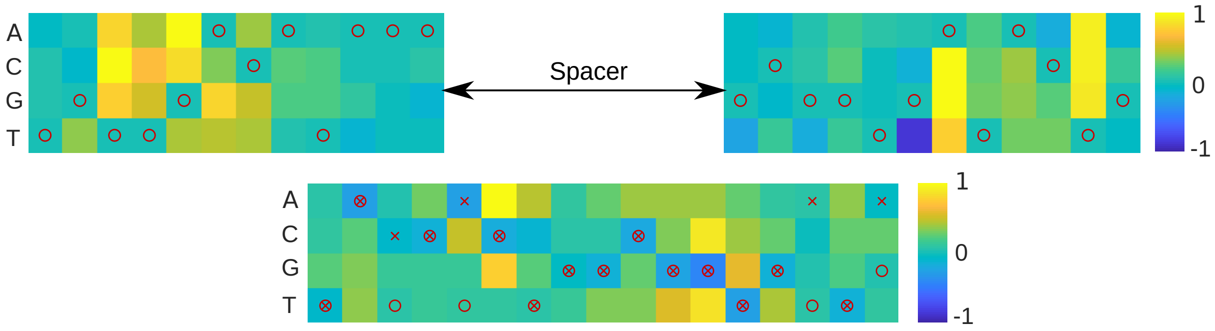


**Figure S1. Energy matrices of RNAP and CI**. We used previously published EMs of RNAp (top) (Lagator *et al.* 2022) and CI (bottom) (Igler *et al.* 2018). In the case of RNAP, the EM included additional positions outside the canonical -10 and -35 binding feet. Red circles mark the Lambda $P_R$ wild-type binding sites for RNAP and $O_{R2}$, while red 'x' marks the wild type $O_{R1}$ sequence. The unit scale is normalized to be between -10 and 1, determined by the highest energy value in each matrix. Methods section *Obtaining the parameters for the model* explains how these relative values were transformed into real energy units $k_B T$.

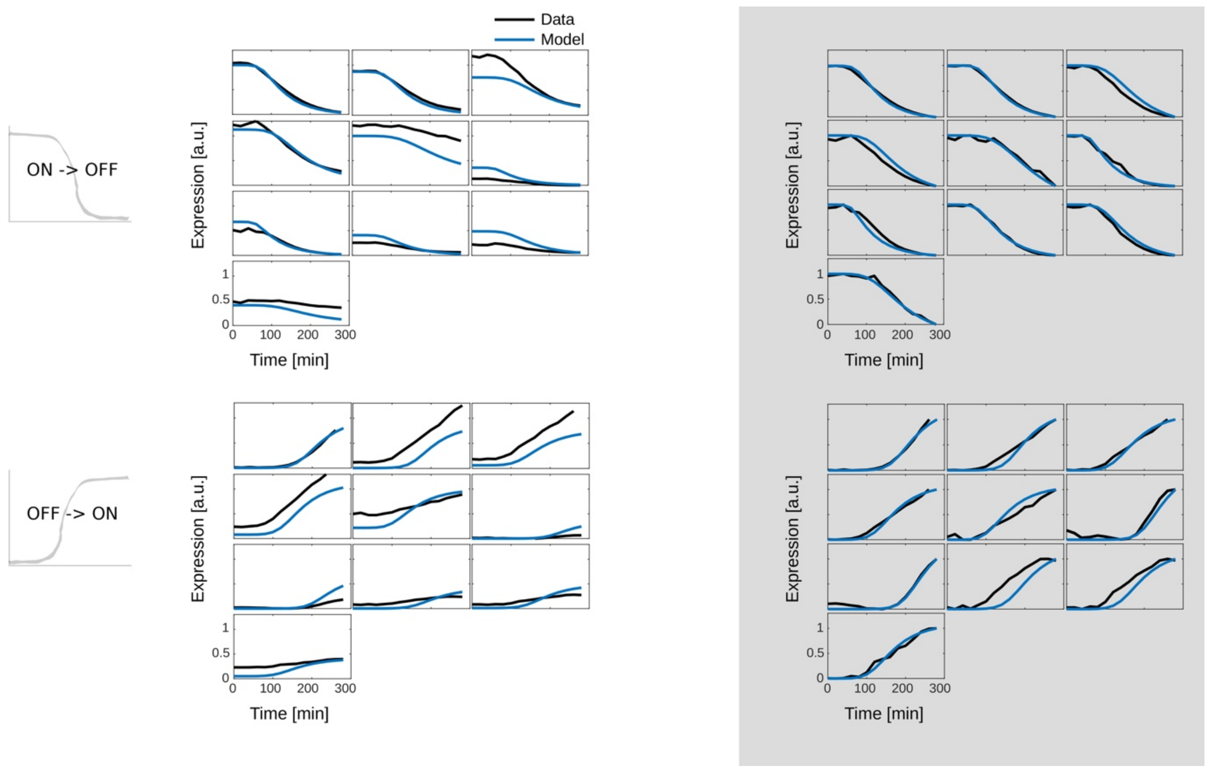


**Figure S2: model to data fit for all tested mutants**. Predictions of gene expression ON->OFF and OFF->ON dynamics for the wild-type (top left of each set of plots) and the nine tested mutants. Plots with white background are based on predictions of all components of the model (I-III in Fig.2B). Gray background shows plots where the thermodynamic component (I and II) was fitted from data, and the mass action kinetics component (III) was predicted.

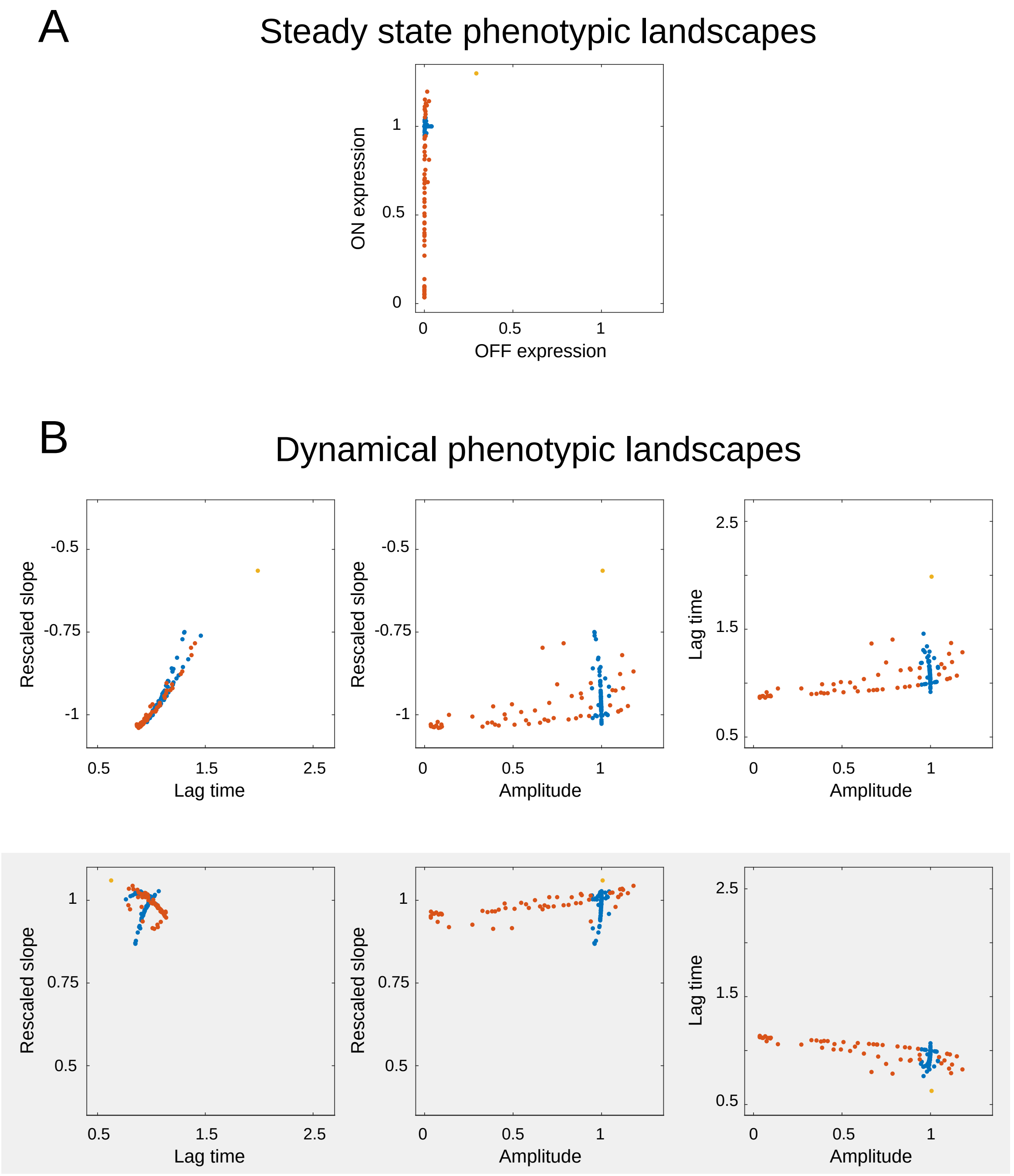


**Figure S3. Phenotypic constraints explored through single point mutations**. A) Phenotypic effects of all possible single point mutations in the $P_R$ promoter on steady-state ON and OFF expression levels. Colors of points indicate whether the mutation affects RNAP binding, CI binding, both or neither – same as in Fig.3. B) Phenotypic effects of all single point mutations on all gene expression phenotypes, shown as two-dimensional interactions between pairs of phenotypes. Top three panels (white background) are the phenotypes of the ON->OFF dynamics, while the bottom three (gray background) are the phenotypes of OFF->ON dynamics. Amplitude represents ON-OFF expression levels. As slope depends on amplitude, we used rescaled slope (slope/amplitude) as the phenotype we report. All units are in the wild-type units, with the exception of OFF expression, which is in the units of ON expression.

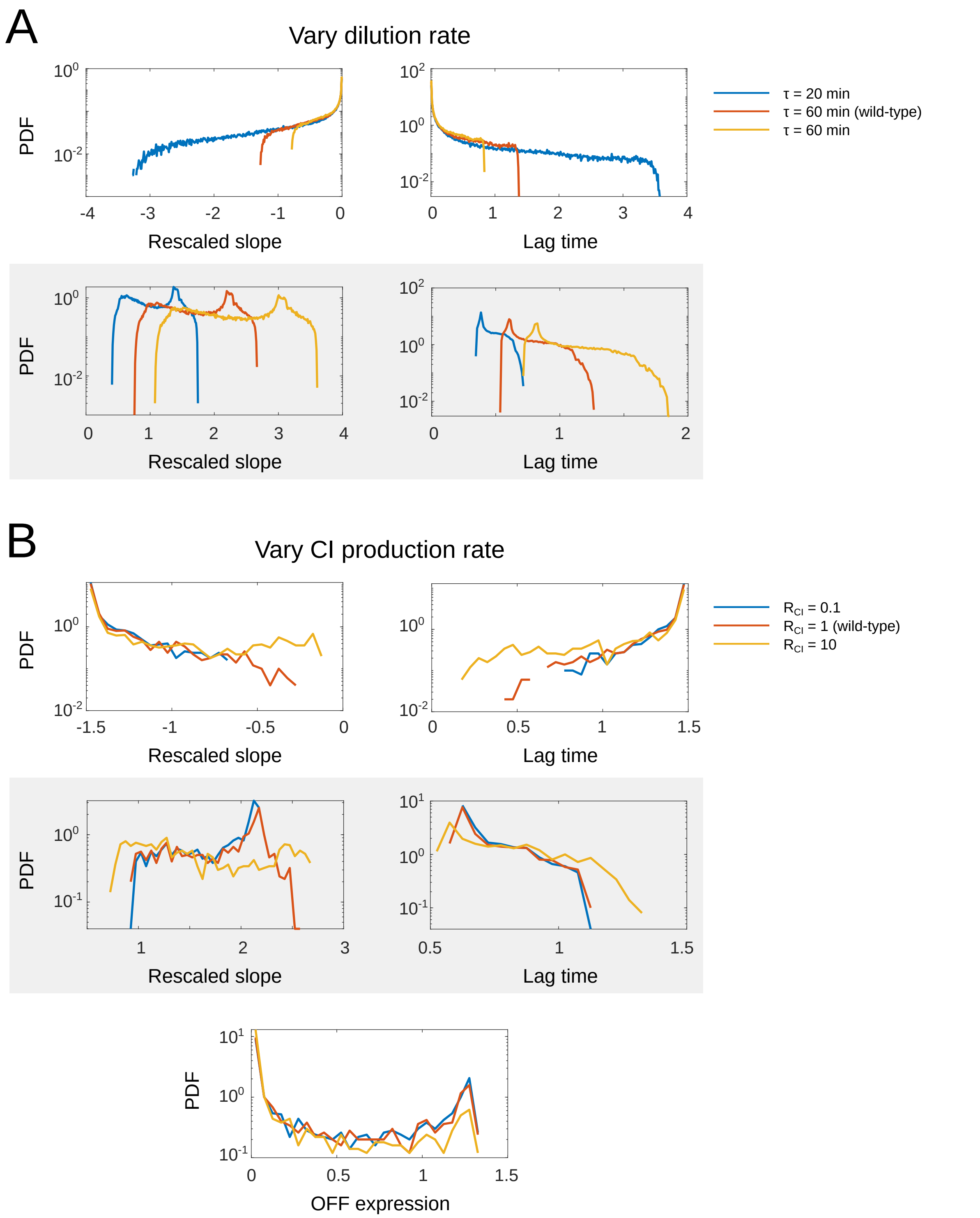


**Figure S4. Probability density function (PDF) of phenotypic values** as a function of A) dilution rate ($\tau$); B) CI production rate ($R_{CI}$). PDFs are shown for all phenotypes that are affected by that parameter (Fig.4A), with white background indicating phenotypes during the ON->OFF dynamics, and gray background indicating OFF->ON dynamics.

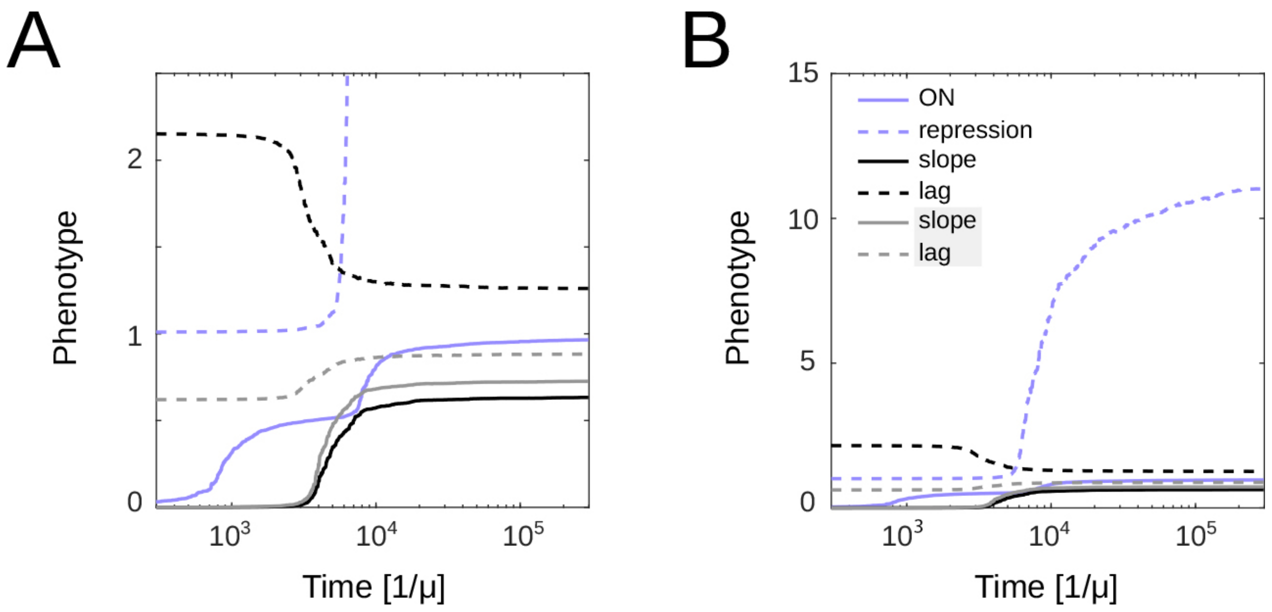


**Figure S5. Non-normalized time trajectories of each phenotype during simulated promoter evolution.** To better represent the evolution of repressible promoters, we show repression (difference between ON and OFF states) rather than OFF expression. While A) focuses on majority of phenotypes, B) shows the zoomed-out version of the same plot to show how repression changes relative to other phenotypes. Units of each phenotype are in their respective wild-type units, with the exception of repression which is in the units of wild-type ON expression. Time units are inverse of mutation rate, μ.

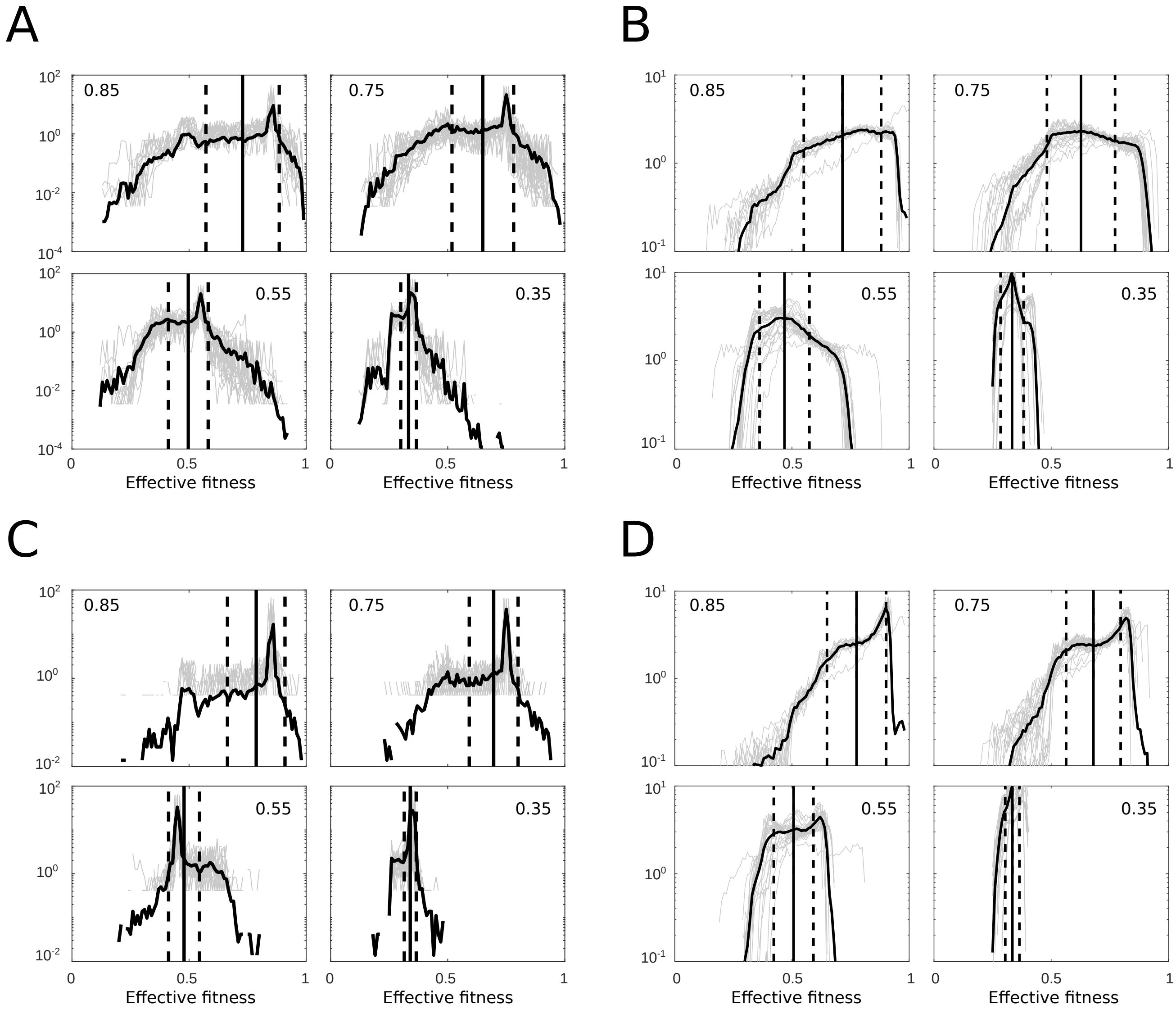


**Figure S6. Comparison of distributions of fitness effects (DFEs) as a function of current fitness, between the full model and the geometric model.** DFEs are shown by simulating all possible double and single mutants, for 30 random sequences with a given fitness (indicated by the number in each plot). A) Double mutants using the full model (note that the same plot is shown in Fig.6A); B) Double mutants using geometric model; C) Single mutants using full model; D) Single mutants using geometric model. Vertical black and dashed lines represent mean and standard deviation of the mean DFE.

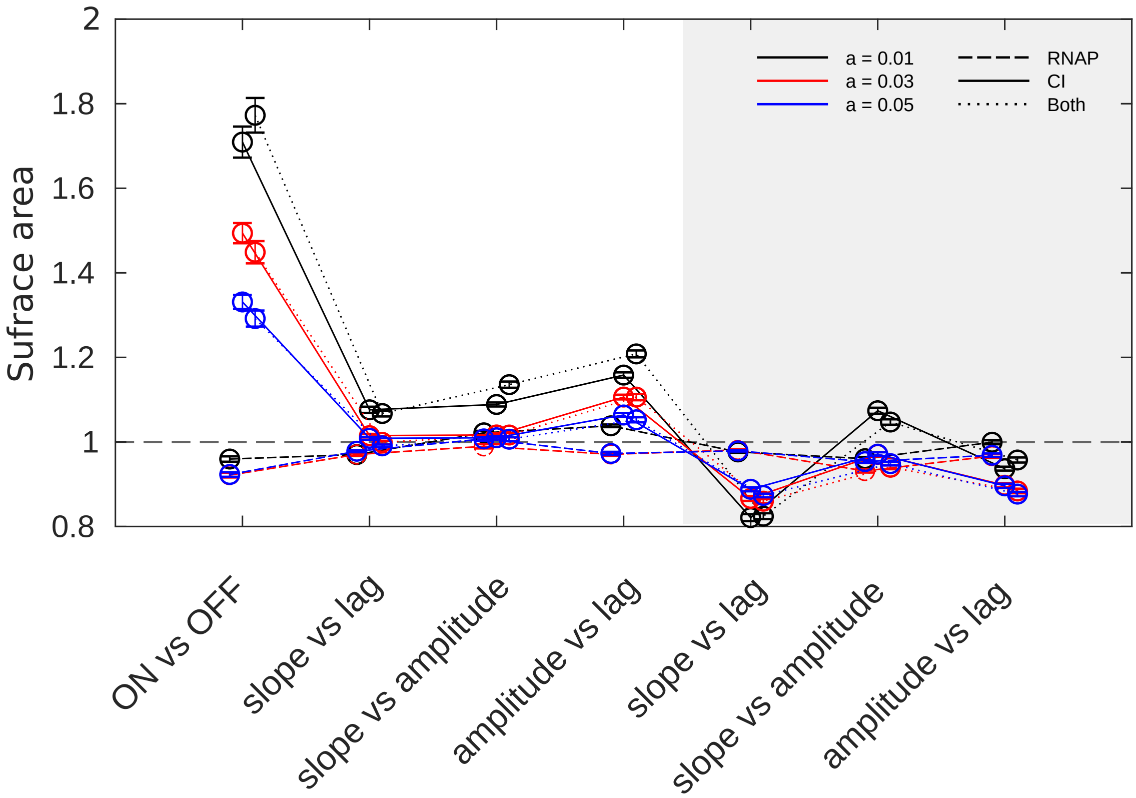


**Figure S7. Normalized phenotypic volume as the function of square size, *a*.** We quantify the normalized phenotypic volume as one of two measures of constraints (see Methods section **Computing the surface area of phenotypic landscapes**). To do this, we consider that each mutation occupies a square in the two-dimensional phenotypic space. Here we show that the estimates of normalized phenotypic surface area did not depend on different sizes of the squares, *a*. We are considering surface area here instead of volume as it was easier to evaluate the effects of *a* on pairs of two-dimensional projections as opposed to a six-dimensional space. White background indicates pairs of phenotypes during the ON->OFF dynamics, while gray background indicates OFF->ON dynamics.
